# Resting networks and personality predict attack speed in social spiders

**DOI:** 10.1101/591453

**Authors:** Edmund R. Hunt, Brian Mi, Rediet Geremew, Camila Fernandez, Brandyn M. Wong, Jonathan N. Pruitt, Noa Pinter-Wollman

## Abstract

Groups of social predators capture large prey items collectively, and their social interaction patterns may impact how quickly they can respond to time-sensitive predation opportunities. We investigated whether various organizational levels of resting interactions (individual, sub-group, group), observed at different intervals leading up to a collective prey attack, impacted the predation speed of colonies of the social spider *Stegodyphus dumicola*. We found that in adult spiders overall group connectivity (average degree) increased group attack speed. However, this effect was detected only immediately before the predation event; connectivity two and four days before prey capture had little impact on the collective dynamics. Significantly, lower social proximity of the group’s boldest individual to other group members (closeness centrality) immediately prior and two days before prey capture was associated with faster attack speeds. These results suggest that for adult spiders, the long-lasting effects of the boldest individual on the group’s attack dynamics are mediated by its role in the social network, and not only by its boldness. This suggests that behavioural traits and social network relationships should be considered together when defining keystone individuals in some contexts. By contrast, for subadult spiders, while the group maximum boldness was negatively correlated with latency to attack, no significant resting network predictors of latency to attack were found. Thus, separate behavioural mechanisms might play distinctive roles in determining collective outcomes at different developmental stages, timescales, and levels of social organization.

**Significance statement:** Certain animals in a group, such as leaders, may have a more important role than other group members in determining their collective behavior. Often these individuals are defined by their behavioral attributes, for example, being bolder than others. We show that in social spiders both the behavioral traits of the influential individual, and its interactions with other group members, shape its role in affecting how quickly the group collectively attacks prey.

## Introduction

Group living can benefit group members through access to mates, protection from predators, and increased foraging opportunities (Krause and Ruxton 2002). A variety of animals engage in cooperative hunting to capture prey that is larger than what they could capture alone. Examples of cooperative hunting can be seen in chimpanzees (Boesch 2002), lions (Stander 1992), wild dogs (Creel and Creel 1995), hawks (Bednarz 1988), killer whales (Pitman and Durban 2012) and invertebrates such as ants (Witte et al. 2010), and social spiders (Whitehouse and Lubin 1999). During collective prey capture, individuals often coordinate their behaviour through social interactions, to maximize their capture success (Stander 1992; Boesch 2002; Pitman and Durban 2012). A shorter latency to attack can reduce the time and effort needed to capture prey, and increases the probability of success, thus conferring important fitness benefits to all group members (Pasquet and Krafft 1992). In addition to coordination through social interactions, groups often rely on particular individuals to expedite collective dynamics (Modlmeier et al. 2014b).

Social network analysis provides tools to quantify interaction patterns and has been instrumental in understanding the dynamics and outcomes of interactions within groups (Wey et al. 2008; Kurvers et al. 2014; Pinter-Wollman et al. 2014; Krause et al. 2015) and in predicting group success (Royle et al. 2012). Different network measures can describe interactions occurring at various organizational levels, such as an individual’s direct interactions with its neighbours, links within a subgroup, or interactions at the whole group level (Lusseau and Newman 2004; Wittemyer et al. 2005; Krause et al. 2007; Wey et al. 2008). The social structure of animal groups often changes over time, and interaction patterns occurring in one period can impact the group later (Blonder et al. 2012; Pinter-Wollman et al. 2014; Krause et al. 2015). For example, social connectivity early in life can predict male mating success several years later in long-tailed manakins (McDonald 2007), social connections to relatives can persist for longer than relationships with non-kin in spotted hyenas (Holekamp et al. 2012), and social stability of subgroups can be maintained over years in sparrows (Shizuka et al. 2014). Thus, temporal dynamics of interactions may occur at different rates at different organization levels (Blonder et al. 2012).

Animals within a society often differ from one another in their behaviour, and these differences can be consistent over time, a phenomenon that has been referred to as ‘personality’ (Sih et al. 2004; Bell et al. 2009; Jandt et al. 2014). Behavioural variation among individuals in a social group can have a considerable impact on group functions (Pinter-Wollman 2012). Only a small amount of variation among individuals may be necessary to have large impacts on the entire group. In the most extreme situations just one ‘keystone’ individual, such as a leader or a tutor, may have a disproportionate impact on the group (Conradt and Roper 2003; Modlmeier et al. 2014b). A keystone can be either a particular individual or a role that different individuals assume at different times (Modlmeier et al. 2014b). Keystone individuals can have an increased interaction rate with other group members (Lloyd-Smith et al. 2005) and their behavioural tendencies can influence interaction patterns (Pike et al. 2008; Sih et al. 2009; Krause et al. 2010; Pinter-Wollman et al. 2011; Firth et al. 2015), thus in turn impacting collective actions (Bansal et al. 2007; Brown and Irving 2014). It is still unknown how keystone individuals influence the performance of a group. Generally speaking, keystone individuals can either perform the work itself or catalyse the work of other group members (Robson and Traniello 1999), for example through social interactions.

Individual differences that are consistent over short time frames may change over longer periods (Stamps and Groothuis 2010). Such changes can alter the relationship between a group’s personality composition and collective outcomes over time. For example, in social insects the task that each individual performs can change with age (Seeley 1982; Tripet and Nonacs 2004) potentially altering the distribution of task performance in the colony. Changes to an animal’s personality may arise from changes in physiological processes such as growth (Biro and Stamps 2008), metamorphosis (Hedrick and Kortet 2012; Wilson and Krause 2012), or sexual maturation (Stamps and Groothuis 2010). Personality may also develop due to changes in the external physical or social environment over time, i.e. experiential factors (Stamps and Groothuis 2010). For example, as group members become familiar with one another, variation among individuals increases and variation within an individual decreases, thus increasing behavioural repeatability (Laskowski and Pruitt 2014; Modlmeier et al. 2014c; Laskowski et al. 2016).

In the social spider *Stegodyphus dumicola* (Araneae, Eresidae) individuals vary in their boldness, and the boldest individual in the colony (referred to as the keystone) affects foraging intensity (attack speed and number of attackers) and mass gain of the entire group (Keiser and Pruitt 2014; Pruitt and Keiser 2014). It is not known how the influence of keystone individuals is imparted in this species, only that keystones have long lasting effects, and that the duration of their impact is proportional to the tenure of the keystone in the group (Pruitt and Pinter-Wollman 2015). A recent model (Pinter-Wollman et al. 2016) predicts that when the boldness of group members is persistent (i.e., is a stable personality trait), social interactions should play a larger role than boldness in shaping collective outcomes. This prediction emerges because who interacts with whom would change more rapidly than boldness if it was highly stable, and boldness does not necessarily determine who interacts with whom (Hunt et al. 2018). In contrast, if boldness is plastic, social interactions may play a smaller role in determining collective outcomes because changes in the collective outcomes can emerge from changes in boldness. Younger animals are often more behaviourally plastic than adults (Scott 1962). Thus, by comparing the social behaviour of both subadult, or juvenile, and adult organisms, we can test when social interactions and when behavioural traits, i.e. boldness, have a larger impact on collective outcomes (Pinter-Wollman et al., 2016).

*S. dumicola* spiders live in colonies of up to several hundred individuals of the same age that exhibit cooperative behaviours such as prey capture and allo-maternal care (Bilde et al. 2007; Junghanns et al. 2017). Enhanced foraging success is thought to be a primary driver of sociality in social spiders (Whitehouse and Lubin 2005), to enable subduing of large prey items (Guevara et al. 2011; Harwood and Avilés 2013). More frequent co-feeding interactions of the same prey item has been observed in sibling groups compared with non-sibling groups in group-foraging subsocial spiders, suggesting that social network structure may play a role in the evolution of social behaviour in spiders (Ruch et al. 2015). Colonies composed of bolder spiders attack more rapidly (Keiser et al. 2014; Keiser and Pruitt 2014; Pruitt and Keiser 2014) and with more individuals (Grinsted et al. 2013; Keiser and Pruitt 2014; Laskowski and Pruitt 2014; Pruitt and Keiser 2014) than colonies with shy individuals.

Here we use social network analysis to determine the temporal scale and the social organization level at which interactions between group members have the most impact on collective prey capture dynamics. We evaluate if the keystone individual influences the group through its role in the group’s social network. We also consider whether these social interactions have long-lasting effects on prey capture success, or if the impact of interactions is immediate and ephemeral. Furthermore, we examine if developmental stage (subadults vs. adults) affects the relationships between personality, social interactions, and collective prey attack. Comparing subadults, with more plastic, emergent personalities, to adults, with more established personalities, allows us to test the predictions of the model of keystone influence detailed above (Pinter-Wollman et al. 2016).

In sum, we consider whether the speed at which *S. dumicola* colonies of either subadult or adult spiders collectively attack prey depends on: *(i)* interactions occurring at different levels of social organization; *(ii)* temporal changes in social structure; and *(iii)* the behavioural and social attributes of the keystone individual.

## Methods

### Animal collection and maintenance

Colonies of *S. dumicola* were collected from roadside *Acacia* trees in the Northern Cape of South Africa in November 2015 (subadults) and March 2016 (adults), transported to the laboratory, and fed crickets *ad lib*. The size of collected colonies ranged between 70-300 individuals and contained only females - males are short-lived and rare (12%) in natural colonies (Henschel et al. 1995). We created 15 groups of 26-30 sub-adult female spiders, from 4 source colonies of subadults, and 24 groups of 10 adult female spiders each, from 3 other source colonies. Individuals from different source colonies were not mixed. Group sizes were larger for subadults because of the small size of those individuals, and because it potentially requires more small individuals to execute a successful attack on large prey (see supplementary Figure S2 for differences in sizes between adults and subadults). The behavioural composition of these groups is detailed below in the ‘*Group composition*’ section. Groups were housed in large round containers (18cm diameter, 8cm depth for subadults and 11cm diameter, 10cm depth for adults) with vertical wire meshes (two 9×6cm sheets positioned 10cm apart for subadults and a 5×5cm sheet for adults) to allow the spiders to build both a retreat and a capture web. Trials were conducted from January until August, 2016.

### Experimental procedure

To determine the effect of interaction patterns at different time scales on prey attack we observed groups over time. Each group was observed for 6.5 weeks. Boldness and prey capture were measured once a week and resting interactions, as detailed below, were observed three times a week with 2-3 days separating each observation. The first resting network was obtained immediately before measuring boldness on Day 4, numbered as four days before measuring prey attack. The second resting network was observed on Day 2, two days after measuring boldness and two days before measuring prey attack. The third resting network was observed immediately before testing prey attack speed, on Day 0. This spacing of measures allowed ample time for the spiders to recover from disturbances due to removing them from their web to determine boldness. Each group was fed with a single 4-week-old cricket once a week, which provides *ad lib* food, after the prey assay (described below), hence all colonies had an equal opportunity to consume prey, and were at a similar state of hunger. We obtained 7 boldness measures for each individual spider, 6 collective prey capture response measures for each group, and 18 resting networks for each group (19 including a final boldness/network observation not used here). We compared the predictive power of the resting networks observed four days, two days, and immediately before each prey capture trial, for explaining the speed of prey attack. This allows us to differentiate between short term (immediately before prey attack), medium (two days), and long (four days) term influences of spider interaction networks.

### Boldness

To determine individuals’ boldness, each spider was tested once a week using an established assay that measured the recovery of a spider from exposure to air puffs, which mimic the approach of an avian predator (Riechert and Hedrick 1993). Spiders react to the air puffs by huddling and remaining still. The faster the spiders resume movement after this simulated threat, i.e., move one body length away from where they were huddled, the bolder they are considered. Bolder spiders tend to participate more in collective prey attacks than shy individuals (Lichtenstein et al. 2017). Boldness is a repeatable behaviour in this species when spiders are kept isolated over days, with a repeatability of 0.63 (Keiser et al. 2014). However, in a social context, boldness is much more plastic and changes as a function of the boldness of the individuals with whom one interacts and over time (Hunt et al. 2018) and may be related with metabolic rate (Lichtenstein et al., 2017). To test boldness, spiders were placed individually in a plastic container (15×15cm) and after 30s of acclimation, two puffs of air were administered to the anterior prosoma using an infant nose-cleaning bulb. Boldness was measured as the latency to resume movement and move one body length. Because bolder individuals resume movement faster, we subtracted the time to resume movement from the maximum duration of the procedure (600s) to create a metric that increases with boldness. We designated as ‘shy’ individuals those with a latency to resume movement of 400-600s, while ‘bold’ individuals were those with a latency to resume movement of 0-200s. The abdomen of each spider was marked uniquely with acrylic paint to track their behaviour over time (Figure 1). We examined how many individuals occupied the role of keystone (boldest) in each experimental group over the 7 weeks of the experiment – this ranged from 1 to 7 individuals (same individual throughout, or a different individual each week). A priori, turnover is expected to be higher for the subadults because they have larger group sizes and therefore more individuals that might replace the keystone individual. We compare the keystone turnover for the two developmental stages using a Wilcoxon rank sum (Mann-Whitney) test. We used linear interpolation to obtain an estimate of boldness on Days 0 and 2 and identify a putative boldest individual on those days, which were in between the weekly boldness measurement taken on Day 4.

**Fig. 1.**
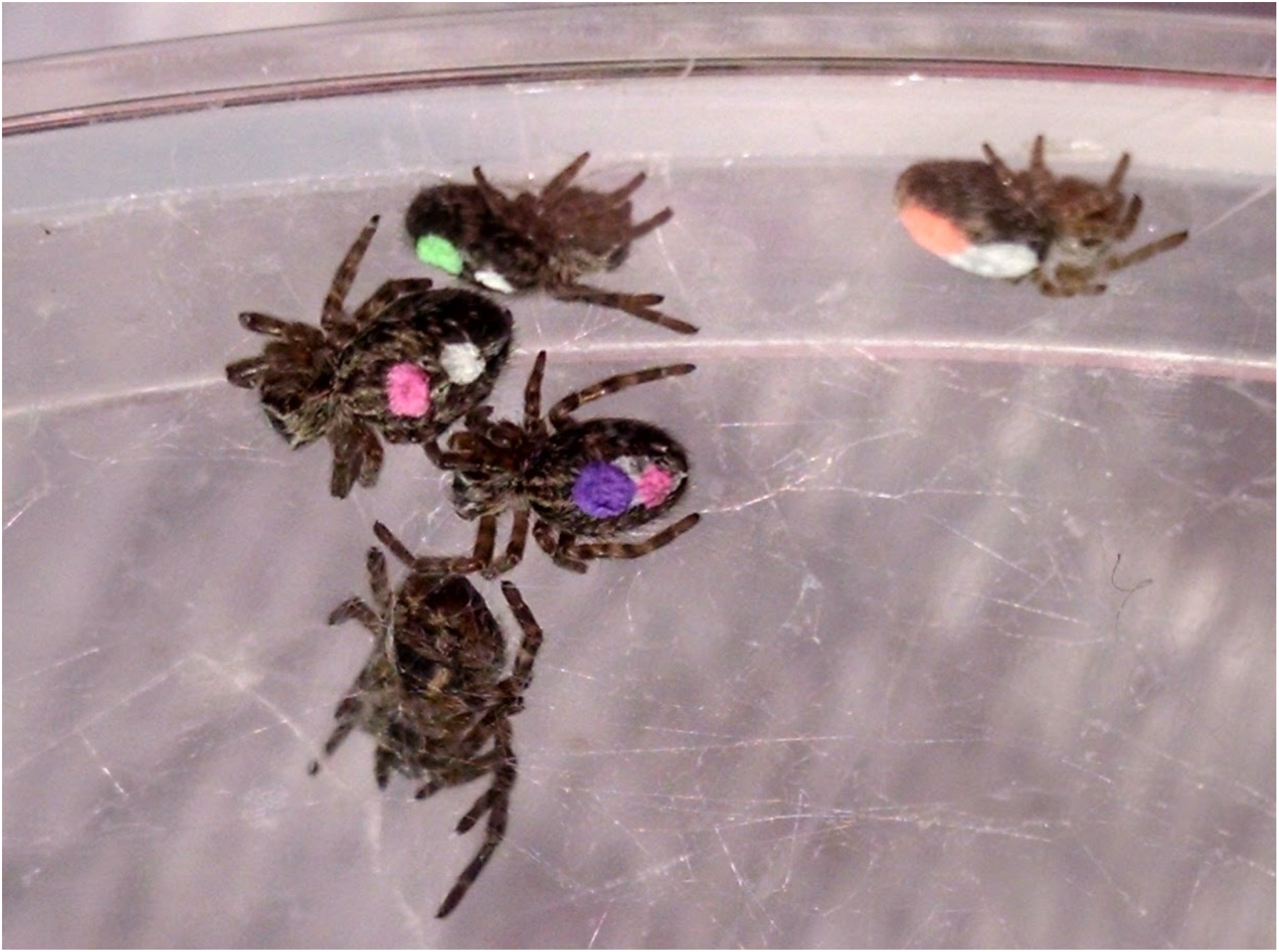
Close-up photograph of resting spiders in the first week of the experiment. The abdomen of each spider is marked uniquely with acrylic paint to track their behaviour over time. The resting network here corresponds to one connected chain (left) and one unconnected node (right).

**Fig. 2.**
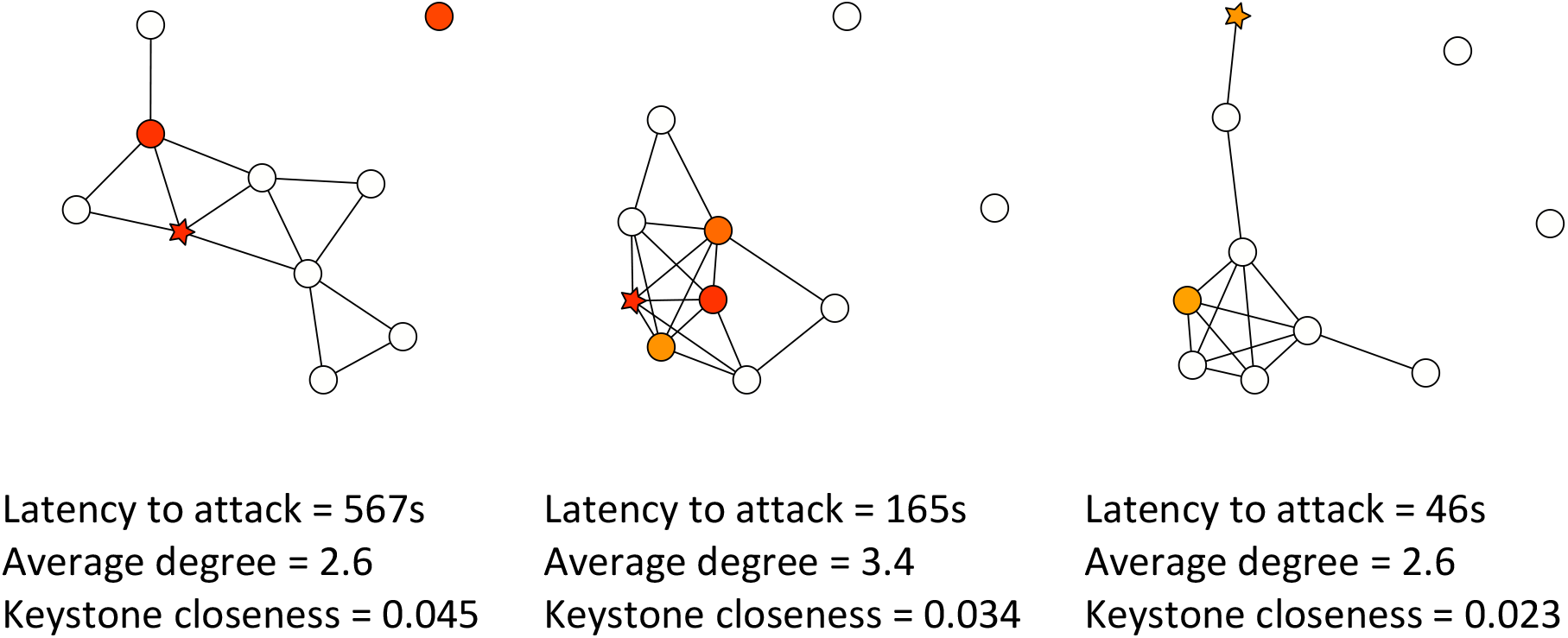
Interaction networks of three sample adult spider groups, immediately before prey attack speed was examined. Each node represents an individual spider and colour represents boldness — redder indicates higher boldness, with the boldest individual marked as a star. Lower keystone closeness and higher average degree are associated with faster latency to attack. Latencies to attack, average degree, and keystone closeness are noted below each network.

### Group composition

To examine the effect of group composition on collective behaviour we assigned spiders to one of three group compositions: all shy, all bold, and ‘keystone’ (all shy individuals plus one bold individual). For subadults, we established five groups of each composition and for adults we established five groups of all bold, nine of all shy, and ten keystone groups. Individuals that were not assigned to experimental groups, including those with a boldness score of 200-400, were returned to their source colony. After the first week of our study, the average boldness of all group compositions converged, with the ‘all bold’ groups reducing their average boldness substantially and the other two group types increasing their average group boldness slightly (Figure S1 A, B). Boldness during week 1 was the boldness recorded before creating the experimental groups, thus before individuals interacted with one another. We used a one-way ANOVA to compare mean group boldness of the three different group compositions, for the latter five weeks of our experiment, excluding week one. Because we did not detect a significant difference in average boldness between the three compositions for the latter five weeks (subadults: F_2,71_=0.739, p=0.48; adults: F_2,117_=2.076, p=0.13), we excluded week one and pooled the remaining data across compositions in all analyses (see also distribution analysis in Table S1). Because boldness was found to be more plastic in a social context compared with isolation, and because the artificially manipulated boldness distributions were quickly returned to their natural skewed state by the spiders’ collective boldness dynamics, we did not find evidence that our treatments had any long-term effects. For further information on changes in boldness over time in a social context see Hunt et al. 2018.

### Social interactions

The physical contacts among spiders were manually recorded three times a week: immediately (1-2 hours) before the prey-capture assay, two days prior to the prey capture assay, and four days before to the prey capture assay. Resting interactions were defined as a physical contact between any body parts of two spiders (Figure 1). Interactions were observed during the day, when spiders are resting and inactive, which is their condition most of the time, unless disturbed by a prey in their web or by a destruction of their web that requires maintenance. Care was taken to note each spider in the colony so that all interactions are recorded. These interactions were used to construct social networks and calculate network variables that indicate individual, sub-group, and group level dynamics. The network variables measuring individual level behaviours were keystone degree, keystone closeness, and maximum boldness in the group (i.e., boldness of the keystone); for the sub-group level, modularity; and for the group level, average degree and degree distribution skewness, as detailed in Table 1.

**Table 1.** Network measures examined

| Level of analysis | Network variable | Description |
| --- | --- | --- |
| Individual | Keystone degree | Number of individuals interacting with the boldest individual |
|  | Keystone closeness | The sum of reciprocal shortest paths from the boldest individual to all other nodes. Measures how well integrated an individual is in the overall network. |
|  | Maximum boldness | The boldness of the boldest (keystone) individual in the group for that week (not a network measure). |
| Sub-group | Modularity | The extent to which the network is divided into modules, measured by the ratio between links within a module and links outward to other modules. Modules were defined based on the Optimal Community Structure algorithm (Brandes et al., 2008). |
| Group | Average degree | The average number of individuals that each individual interacts with. Quantifies overall network connectivity. |
|  | Skewness of degree distribution | The skewness of the degree distribution. Larger absolute values indicate that degree is less evenly distributed among individuals in the group. |

To calculate the 2 individual-level network measures (Table 1), we had to first identify which individual was the boldest, i.e. occupying the keystone role each week. In this system, the individual with the highest boldness assumes the keystone role, regardless of its identity (Pinter-Wollman et al. 2017b). Thus, the role of keystone is not necessarily maintained by a specific individual. In a social setting, individuals change their boldness over time (Hunt et al. 2018) and so boldness ranks among individuals may also shift. When more than one spider exhibited the same maximum value, we took the average network value of those individuals (this happened 2 out of 74 times for the subadult spiders and 1 out of 120 times for the adults). When all spiders had zero boldness, we identified the keystone individual as the boldest spider in the previous week (this did not occur for the subadults, and 13 out of 120 times for the adults); where this was not possible, we calculated network values as an average across all individuals in the group that week (7 out of 120 times for the adults).

### Prey response

To determine the speed at which groups attacked prey collectively, we examined the groups’ latency to respond to vibrations on their capture web (Grinsted et al. 2013). We used a custom-made vibratory device assembled from an Arduino Uno board, a vibratory motor, and a metal wire, directed at a 1×1cm piece of paper placed in the capture web (Pinter-Wollman et al. 2017b). The stimulus was always placed on the capture web at the same distance (4cm) from the nest retreat, where most spiders were gathered, to control for any effects the distance of the stimulus might have on the response of the group. The Arduino board was programmed to vibrate the piece of paper in pulses that varied randomly between 0.5-1.5 sec in both the duration of the vibration and the pauses between vibrations, to simulate the irregular vibrations that a prey makes when captured in the web (Hedrick and Riechert 1989). The paper was vibrated until a spider touched it, to avoid habituation to our stimulus, or until 10 minutes elapsed, in which case the trial was stopped (Pinter-Wollman et al. 2017b). As the first individual left the retreat, others followed, creating a collective response. The first individual to leave the retreat was not necessarily the one closest to the simulated prey (personal observations). When no attack took place, we set the latency to attack to ten minutes. We noted the identity of the first individual(s) to touch the stimulus, as well as the identity of all the individuals that left the nest during the attack as participants, so that we could assess whether the keystone (boldest) individual participated in prey attack. Both adult and subadult groups responded to the simulated prey in a similar manner (Figure S3).

### Data analysis

To examine the relationship between social network structure, boldness, and prey attack we used censored mixed regression models. We considered six variables as predictors of latency to attack, as detailed in Table 1. These were included as fixed effects interacting with the effect ‘Day’ which accounted for the number of days before the attack assay (4, 2, or 0). This approach allowed us to determine the timescale on which social interactions act on prey capture. We constructed separate models for adult and subadult spider behaviour, and each model included 5 weeks of data (weeks 2-6, excluding week 1). For the adult spiders these included N=360 resting network observations (24 colonies x 5 weeks x 3 observations per week) and for the subadults N=224 (15 colonies x 5 x 3, except N=74 for Day 0 because of 1 missing network observation). Because latency to attack was right-censored at 600 s, with 76% of subadult trials and 65% of adult trials resulting in an attack, we used censored regression (Tobit) models with the R package ‘censReg’ (Henningsen, 2017). The response variable, latency to attack, was log-transformed to adhere to the model assumption that the error term is normally distributed. To account for variation among groups and source colonies we included group identity as a random effect and source colony identity as a fixed effect. We further included a time-varying residual component as a random effect to account for changes over the 5 weeks. Because we did not have an *a priori* prediction regarding which network variable would best explain collective prey attack, we identified a suitable model for the adults and subadults by first estimating all 63 possible models that linearly combined one or more (i.e., not including interactions) of the 6 fixed effects listed in Table 1. We calculated the Akaike weight of the 63 models (Burnham et al., 2011). These model weights were then used to estimate the relative importance of the six predictor variables under consideration (Table 1). The importance of each predictor is determined by summing the Akaike weights of each model in which it appears (Symonds and Moussalli 2011). If a certain predictor appears in many of the top models, its summed Akaike weight will tend toward 1. On the other hand, if it only appears in the weaker models, its weight will tend toward 0 (Symonds and Moussalli 2011). This procedure results in a ranking of predictor variables in terms of performance, and one can interpret the predictor weight as a probability that it is a component of the best model (Burnham and Anderson 2002; Symonds and Moussalli 2011). We selected the top-ranked predictor variable at the individual level, and the top-ranked predictor at the group or subgroup level, to obtain a parsimonious model for the adult and subadult behaviour with two main predictors in each model (Tables 2, 3, 4). We find this predictor-ranking approach to be preferable to full-model averaging because it allows us to identify a particular model with properly estimated predictor standard errors and without excessive model complexity (full-model averaging results are available in Table S7, S8). We checked for multicollinearity between predictors in the final models by calculating their corrected generalized variance inflation factors (Fox and Monette 1992), using the R package ‘car’ (Fox and Weisberg 2011). There was low collinearity in both the adult model (Table S3) and subadult model (Table S5). To assess whether the keystone spider’s role in the interactions network (closeness and degree) was correlated with its boldness we calculated the Pearson’s correlation between these measures.

**Table 2.** The Akaike weights for the attack speed predictors, as calculated across all possible models (see Tables S3 and S4). The top two predictors are selected for the final models, one individual and one group or subgroup level.

| Level of analysis | Network variable | Predictor weights |  |
| --- | --- | --- | --- |
|  |  | Adults | Subadults |
| Individual | Keystone degree | 0.14 | 0.52 |
|  | Keystone closeness | **0.81** | 0.15 |
|  | Maximum boldness | 0.29 | **0.97** |
| Subgroup | Modularity | 0.36 | 0.42 |
| Group | Average degree | **0.46** | **0.47** |
|  | Skewness of degree distribution | 0.06 | 0.18 |

**Table 3.** Statistics of the selected model for attack speed in adult spiders. *denotes significance at p < 0.05. See Table S2 for information on random effects and t values.

| Level of Analysis | Coefficient | Days before | Estimate log(seconds) | Standard error | p |
| --- | --- | --- | --- | --- | --- |
| Individual | Closeness of keystone | 4 | 10.715 | 6.682 | 0.109 |
|  |  | 2 | 12.766 | 6.269 | **0.042\*** |
|  |  | 0 | 18.842 | 7.205 | **0.0089\*** |
| Group | Average degree | 4 | -0.172 | 0.129 | 0.182 |
|  |  | 2 | -0.239 | 0.150 | 0.111 |
|  |  | 0 | -0.367 | 0.175 | **0.036\*** |

**Table 4.** Statistics of the selected model for attack speed in subadult spiders. *denotes significance at p < 0.05. See Table S4 for information on random effects and t values.

| Level of Analysis | Coefficient | Days before | Estimate log(seconds) | Standard error | p |
| --- | --- | --- | --- | --- | --- |
| Individual | Maximum boldness | 4 | -0.0050 | 0.0015 | **< 0.001\*** |
|  |  | 2 | -0.0049 | 0.0016 | **0.002\*** |
|  |  | 0 | -0.0053 | 0.0016 | **< 0.001\*** |
| Group | Average degree | 4 | 0.005 | 0.0247 | 0.8318 |
|  |  | 2 | -0.041 | 0.0314 | 0.1819 |
|  |  | 0 | -0.025 | 0.0245 | 0.317 |

## Results

The adult spider model included average degree (predictor weight 46%), as the group or subgroup-level fixed effect, and closeness of the keystone individual (predictor weight 81%), as the individual-level fixed effect (Tables 2, 3). The subadult spider model included average degree (predictor weight 47%), as the group or subgroup-level fixed effect, and maximum boldness (predictor weight 97%), as the individual-level fixed effect (Tables 2, 4).

Network measures impacted collective prey attack in groups of adult spiders. Adult spider groups that were overall more connected (high average degree) attacked the prey stimulus more quickly. This relationship was found only for resting networks observed immediately before testing collective prey attack, but not for networks measured earlier (Table 3). At the level of the individual, when the groups’ boldest (keystone) individual was more closely connected to all other individuals (closeness centrality), prey attack was slower. This relationship between prey attack and the closeness centrality of the boldest individual was retained for resting networks obtained two days prior to prey attack (Table 3).

No relationship between the network measure average degree and attack speed was found for the groups of subadult spiders (Tables 4). We also ran an alternative model that included degree of keystone as a second individual-level, network effect (predictor weight 52%) but this was also not significant (Table S6). However, the boldness of the boldest individual in the group, i.e., the keystone’s boldness, significantly reduced latency to attack i.e., groups with a bolder keystone attacked more quickly (Table 4).

In the adult spiders, the boldness of the boldest (keystone) individual was not associated with its degree in the before-prey resting network (Pearson’s correlation: r = 0.039, t = 0.428, df = 118, p = 0.670), or its closeness (Pearson’s correlation: r = −0.007, t = −0.074, df = 118, p-value = 0.941). This was also the case for the subadults’ boldest spiders and their degree (Pearson’s correlation: r = −0.212, t = −1.837, df = 72, p-value = 0.070) and closeness (Pearson’s correlation: r = −0.069, t = −0.5909, df = 72, p = 0.556). There was high turnover in the role of keystone (boldest) individual in both in the adult (4.38 ± 0.68, mean ± standard deviation) and subadult (5.67 ± 0.87) groups (number of different individuals in the role out of a maximum of 7). There was a significant difference between the adult and subadult distributions of keystone turnover (Wilcoxon rank sum test: W=230, p=0.0003).

Keystone individuals participated in 7-29% of prey attacks. In the adult groups on the first week, 24 trials resulted in 21 prey attacks. 17 of these attacking groups had a keystone individual (the remainder of the groups were set up as all-shy individuals), and 5 (29.4%) of the attacks by these groups included the keystone individual. In contrast, on weeks 2-6, 73 of 120 trials resulted in prey attacks. In 70 of these groups there was an individual that had the highest boldness (i.e., was a keystone), and in only 5 (7.1%) of these groups, the keystone individual participated in the prey attack. This decrease in keystone participation in prey attack after week 1 is consistent with previous findings (Figure S2 of Pruitt and Pinter-Wollman 2015). In the subadult groups, on the first week, 15 trials resulted in 12 prey attacks and only in one (8.3%) of these attacks a keystone individual participated. In weeks 2-6, there were 75 trials, 56 resulting in prey attack, 4 (7.1%) of which had keystone participation.

## Discussion

The structure of resting interaction networks measured immediately before prey attack predicted the attack speed of adult social spiders. Higher overall connectivity (average degree), a group-level measure, led to faster attack, though only in the resting interactions measured pre-stimulus. Furthermore, the more connected to others (greater closeness centrality) was the boldest individual (keystone), the slower the collective prey attack speeds, when considering both the networks observed immediately and two days prior to the prey attack. Therefore, the social connectivity of the keystone to the rest of its group, but not its boldness, was found to be significant in groups of adults. The opposite finding was observed for the subadult spiders: groups with a bolder keystone individual attacked the prey stimulus more quickly, but we did not detect any significant associations between subadult resting network measures and attack speeds. These findings support our hypothesis that subadult spiders will rely on behavioural traits likely because younger animals are often more behaviourally plastic than adults (Scott 1962). Our results further support the hypothesis that adult groups rely on social interactions more than on behavioural changes to modify collective dynamics. We also found that the individual-level predictor had a noticeably higher predictor weight compared to the group-level predictor in both the adults and subadults: 81% vs 46% in the adults, and 97% vs 47% in the subadults (Table 3). This seems to confirm the relevance of the ‘keystone individual’ concept in this species.

Influential ‘keystone’ individuals in social groups have a large effect on their social environment relative to their abundance (Modlmeier et al. 2014b). While this influence can be mediated through the behavioural traits of the keystone individual, it may also depend on its social interactions. Here we found that the behavioural traits of the keystone are not the only feature that impacts group success. When the boldest individual in an adult spider group had a lower closeness, i.e., was less connected with other resting spiders, the colony attacked prey more quickly. Thus, we find that the impact of the keystone’s defining trait – high boldness – was mediated by its interactions with the rest of the social group. Other studies define keystone individuals according to their centrality in an interaction network (Lusseau and Newman 2004; Vital and Martins 2013; Modlmeier et al. 2014b) and deem central individuals as critical for the stability of their society. Our findings suggest that *both* the behavioural traits and role in the social network should be considered when defining keystone individuals in some contexts. The boldness of the keystone individuals was not associated with network closeness or degree in either adults or subadults, which suggests that centrality in an interaction network is not merely a direct consequence of individual behavioural characteristics but may result from a different process.

Our results suggest that a keystone individual affects group dynamics either through its own behaviour, or through influencing the formation and dissolution of social interactions. We found low direct participation of keystone individuals in prey attacks (7-29%), which is similar to keystone participation seen in other studies of this system (Pruitt and Pinter-Wollman 2015). Direct keystone participation in prey attack is most common during the first three days after *S. dumicola* colony establishment (Pruitt and Pinter-Wollman 2015) or after a new boldest individual is introduced (Pinter-Wollman et al. 2017b). Keystones rarely participate in prey capture directly in established colonies (Pruitt and Keiser 2014). Seismic recruitment signals have been indicated in *S. sarasinorum* (Bradoo 1980) and other spider genera such as *Theridion saxatile* (Norgaard 1956). Likewise, adult *S. dumicola* are observed to catalyse foraging participation by juvenile spiders without becoming directly involved (Modlmeier et al. 2015). Thus, faster attack speeds may result from keystone individuals signalling information to the rest of the group about the presence of prey through, for example, vibrations on the web. Such signalling does not necessarily require physical proximity, only connection through the capture web, thus it is consistent with the peripheral position of the keystone individual in the proximity networks. In addition, the observed fast attacks in groups with high connectivity that we observed could point to another mechanism underlying the effects of interactions on behaviour. Recent work shows that in this study system boldness is impacted by proximity interactions (Hunt et al. 2018). Thus, it is possible that individuals in groups with high connectivity have more opportunities to modify each other’s behaviour and shape overall group boldness composition, which impacts prey attack.

Interaction patterns had effects on collective outcomes depending on when they occurred. The closeness of the keystone was a significant predictor of attack speed up to two days before the stimulus. This effect duration may be a result of persistence in occupying certain social roles in the interaction network or, as previous research indicates, because keystone spiders have legacy effects on their social environment (Pruitt and Pinter-Wollman 2015). A keystone individual’s long-term influence on its group can result from influencing the behaviour of others or their interactions. For example, in primates the removal of certain individuals who engage in policing behaviour affects the group’s social network structure, even in situations, such as play, in which the removed individuals did not participate (Flack et al. 2006). *S. dumicola* colonies containing bolder keystones attack with more spiders, even after the keystone has been removed, suggesting that keystone individuals shape the overall structure of the social network, and/or change the group members’ individual characteristics (Pruitt and Pinter-Wollman 2015). The lack of relationship between the boldness of the boldest individual in the group and prey attack speed in the adults, could be due to these long-term effects and the dynamics of boldness that we observed. It is possible that it is the boldness of the boldest individual many days before the prey attack that has the largest impact on group dynamics, rather than the boldness of the currently boldest individual. Bold spiders may actively contribute to a process of social niche construction, whereby repeated social interactions reinforce and enhance inter-individual differences in behaviour such as boldness and prey attack speed (Laskowski and Pruitt 2014).

The impact of overall group connectivity (average degree) on the adult spiders’ response to prey suggests that most group members have a role to play in coordinating collective predation, even without needing to participate directly in prey attack. For instance, the presence of fellow colony members may catalyse increased foraging aggressiveness in individuals that are already predisposed to participate in prey capture (e.g., bold individuals (Wright et al. 2015, 2016)). Such catalytic effects have been observed in the context of nightly web repair in *S. dumicola* (Keiser et al. 2016). Furthermore, group level interactions impacted collective prey attack only on a short-time scale. The significant effect of group connectivity (average degree) immediately before, but not two and four days prior to prey capture on its speed, suggests that group level measures are important for immediate dynamics, such as the coordination of motion during prey capture (Krafft and Pasquet 1991). For example, it is possible that spatial proximity between individuals facilitates more rapid information through vibrations or allows individuals to better distinguish vibrations from prey and vibrations from colony mates. Such immediate effects of social connectivity are consistent with observations that small and confined artificial nest scaffolds facilitate rapid prey capture (Modlmeier et al. 2014a). Tighter proximity among group members may increase the frequency of social interactions among all group members (Modlmeier et al. 2014a). Future work on the impact of nest architecture on social interactions could uncover a potential mechanism underlying social interaction patterns (Pinter-Wollman et al. 2017a).

If individuals are behaviourally flexible, the group may be less reliant on social interactions to coordinate activities. Subadult spider groups attacked the prey stimulus more quickly if they had a bolder keystone individual, however, no association was found between their resting networks and attack speeds. These findings are consistent with previous work showing that groups with bolder keystones attack prey faster and with more attackers (Keiser and Pruitt 2014). The lower importance of social network structure in subadults compared with adult spider groups could be related to greater behavioural plasticity in the subadults. In a new social environment *Stegodyphus* spiders show higher within-individual variation in boldness before becoming more behaviourally consistent with time (Laskowski and Pruitt 2014; Modlmeier et al. 2014c; Laskowski et al. 2016). Earlier developmental stages show more behavioural flexibility in a variety of animal systems (Stamps and Groothuis 2010). Here we found high turnover in keystone identity in both adults and subadults, with higher turnover in the subadults. Because adult spiders are perhaps more constrained in adjusting their behavioural traits, they may be more reliant on social interactions to achieve a beneficial group-level behavioural phenotype, because who interacts with whom can be changed more easily. Thus, effective group-level hunting behaviour, which is a key function of sociality in social spiders (Whitehouse and Lubin 2005), could depend on a balance between the plasticity of individual-level behavioural traits and the ease of modifying social interactions (Pinter-Wollman et al. 2016), which may vary according to developmental stage.

Finally, group size may have played a role in the differences we observed between adult and subadult groups. In our experiments, adult groups had 10 individuals and subadults 26-30 and so development stage and group size are not independent of each other in this study. Individuals in smaller groups have been found to participate more in collective prey capture, probably as a consequence of necessity rather than a change in behaviour (Wright et al. 2015). Thus, it is possible that, if not all individuals are required for prey capture in large groups, interactions among all group members are less important than in small groups. Furthermore, larger groups are more likely to contain skilled or experienced individuals, assuming they have greater diversity in ability or experience; this is referred to as the ‘pool of competence’ hypothesis (Morand-Ferron and Quinn 2011). However, subadults are all likely equally inexperienced and unskilled at hunting and so variation in experience in a large group may not be a suitable explanation for the differences we observed between the adult and subadult groups. Finally, the containers occupied by subadults were somewhat larger than those of the adults (to accommodate the larger number of individuals). However, subadult group members tended to all cluster in one small area and not utilize more space than the adult spiders in their smaller containers. Therefore, we expect group size to have a smaller impact on the differences between subadult and adult groups than the difference in developmental stage.

## Conclusion

Our study shows that social interactions are important in determining the speed of collective predation in social spiders and that the impact of interactions may differ according to developmental stage, time scale, and level of social organization. Developmental stages may vary in the importance of social interactions versus shifts in individuals’ behavioural tendencies (Pinter-Wollman et al. 2016). Furthermore, different types of interactions on different social organizational levels may vary in the duration of their effects. For example, we found that an individual-based network measure had longer lasting effects than a group-level measure. Finally, our results indicate that the social role of keystone individuals within a group can have a longer-term effect on collective hunting success. This finding emphasizes the importance of observing both the social structure of individuals in a group and noting inter-individual differences in personality characteristics, for understanding the success of some animal societies. Although keystone individuals can occupy a wide variety of roles across different taxa (Modlmeier et al. 2014b), our work suggests that a combination of network and behavioural trait characteristics could provide a more precise definition of keystones in many contexts in future work.

## Acknowledgements

We thank the South Africa Department of Tourism, Environment, and Conservation for providing permits for animal collection (FAUNA 1072/2013 and 1691/2015) and Colin Wright and James Lichtenstein for collecting spiders in the field. We further thank Arne Henningsen for guidance on the ‘censReg’ R package.

## Data availability

The datasets generated and analysed during the study will be available in an online data repository upon acceptance of the paper.

## Author contributions

ERH analyzed the data and drafted the manuscript, NPW and JNP designed the study, BM, RG, CF, BW, and NPW collected the data, and all authors contributed to the final version of the manuscript.

