## Supplementary Material for "Resting networks and personality predict attack speed in social spiders"

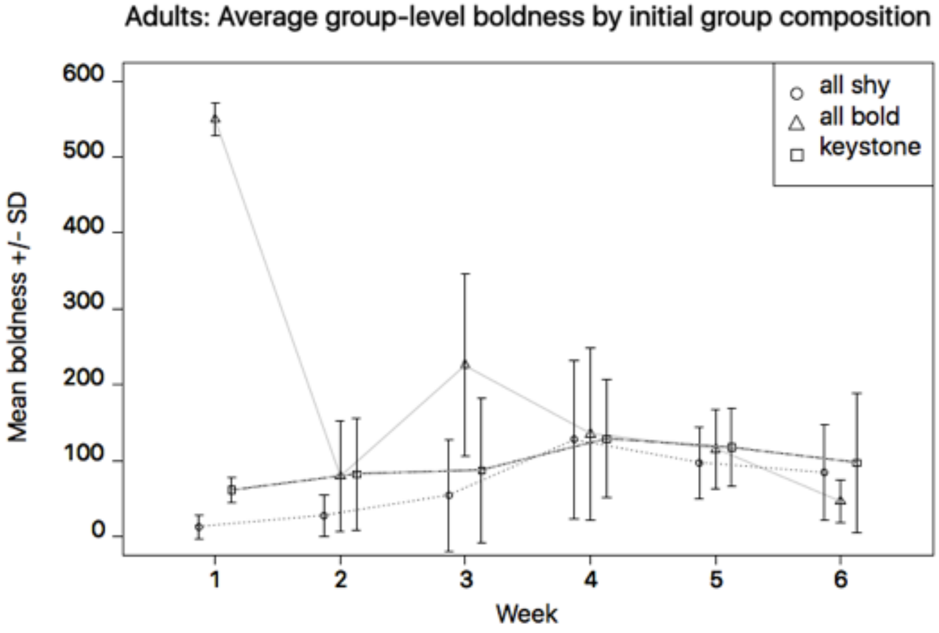


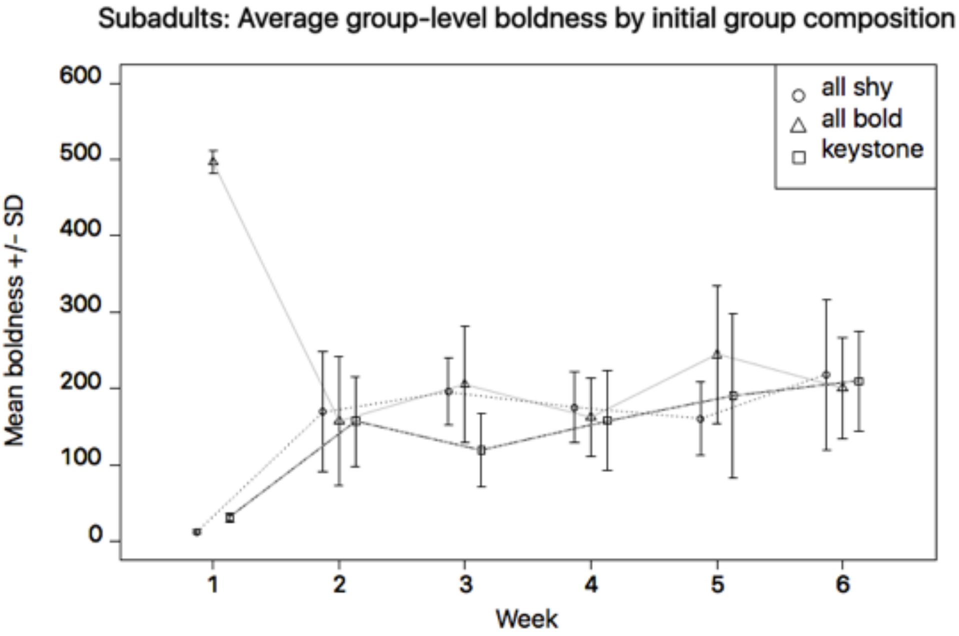


**Figure S1-A.** Mean group boldness by initial boldness composition. Top: adults and bottom: subadults. Error bars indicate standard deviation. For adults, N=9 groups of all shy, N=5 groups of all bold, and N=10 groups of keystone groups. For subadults, N=5 groups for each of the three compositions. For both adults and subadults, mean group boldness converged after the first week.

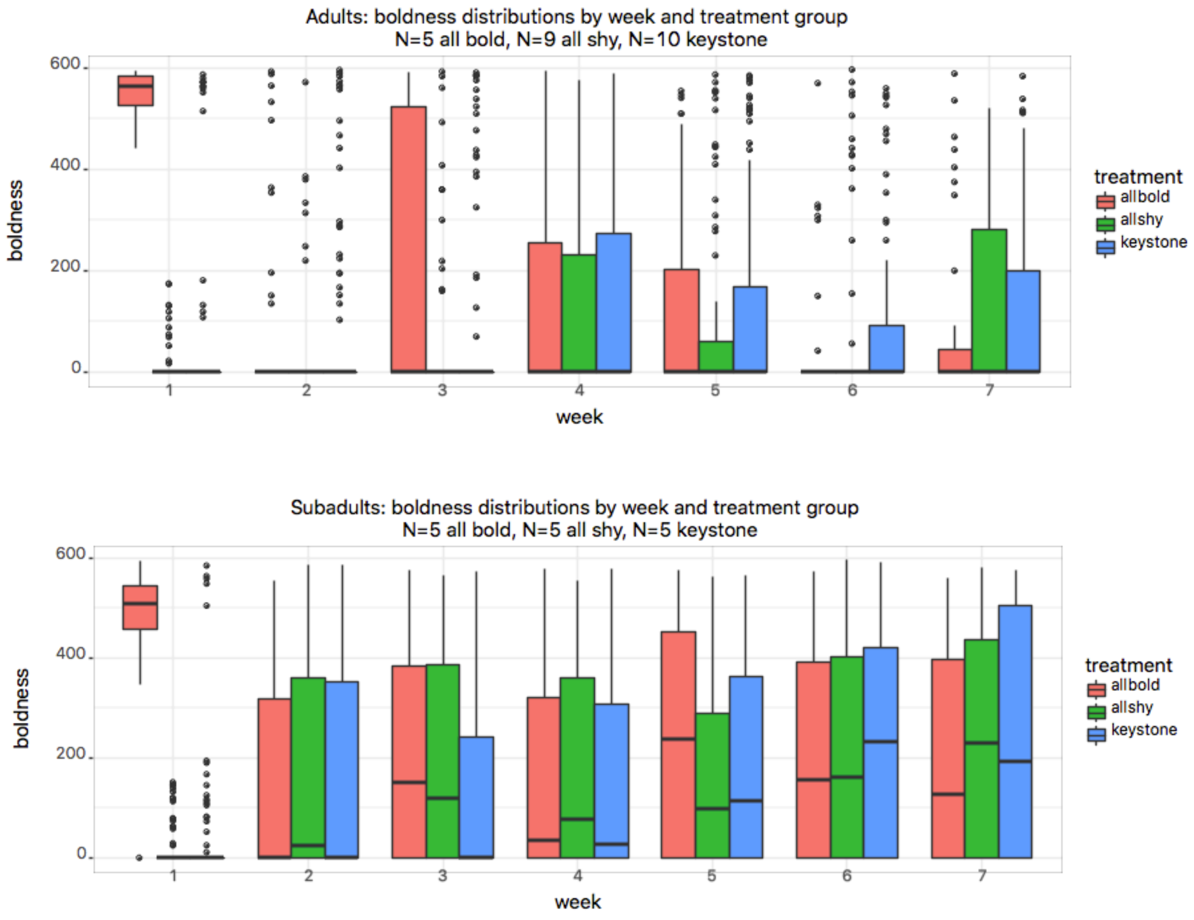


**Figure S1-B.** Boxplots of the adult and subadult spider boldness by week and treatment. The thick line is the median, the lower and upper hinges are the first and third quartiles, the whisker is 1.5*IQR, and the single points are outliers beyond this. The boldness substantially converges in weeks 2-6, justifying the pooling of treatments.

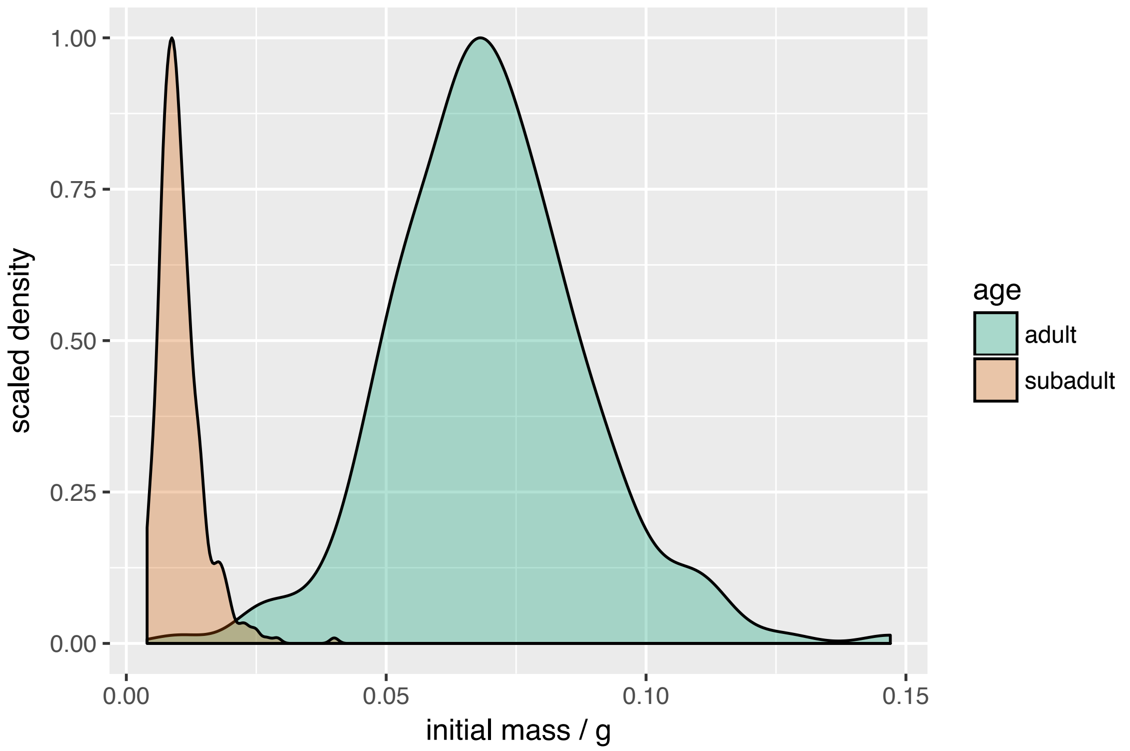


Figure S2. Distribution of initial spider mass for the two developmental stages (subadult or adult) scaled to 1 on the y-axis to allow comparison. Subadults have the physical features of adults but they are much smaller.

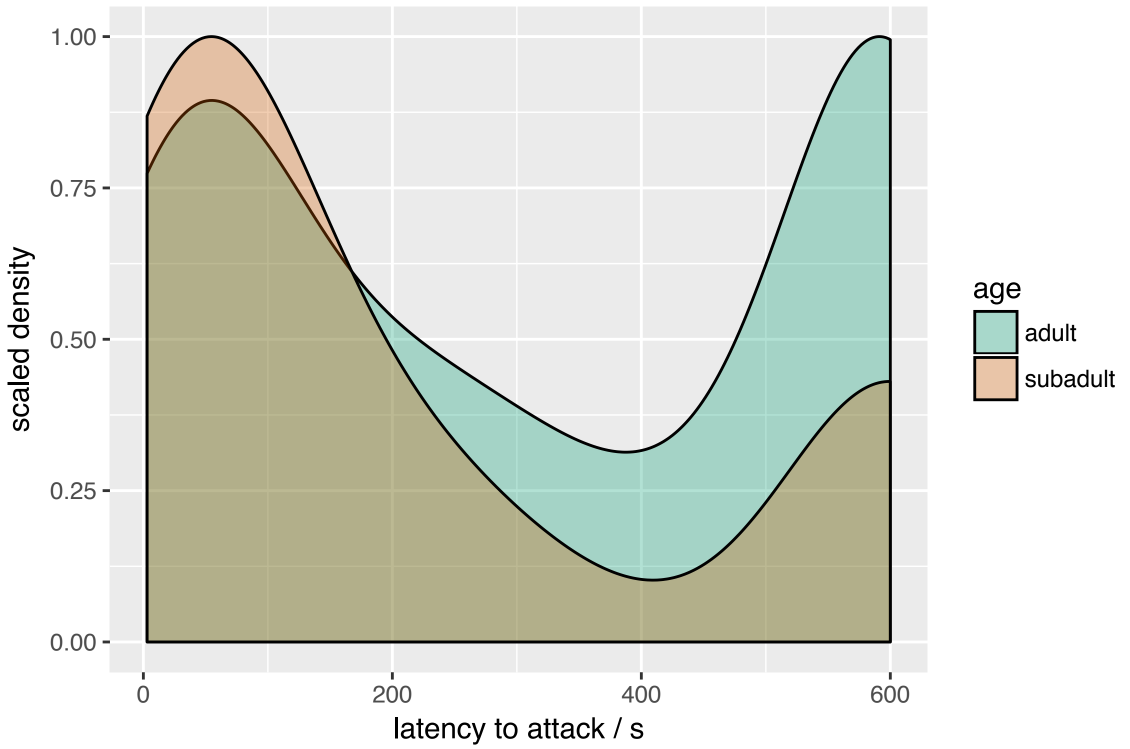


**Figure S3.** Distribution of latency to attack for the two developmental stages (subadult or adult). Scaled to 1 on the y-axis for comparability. There is right-censoring at t=600s, which is the time given for no attack (i.e. > 600s). The two distributions are broadly similar in actual attack speeds.

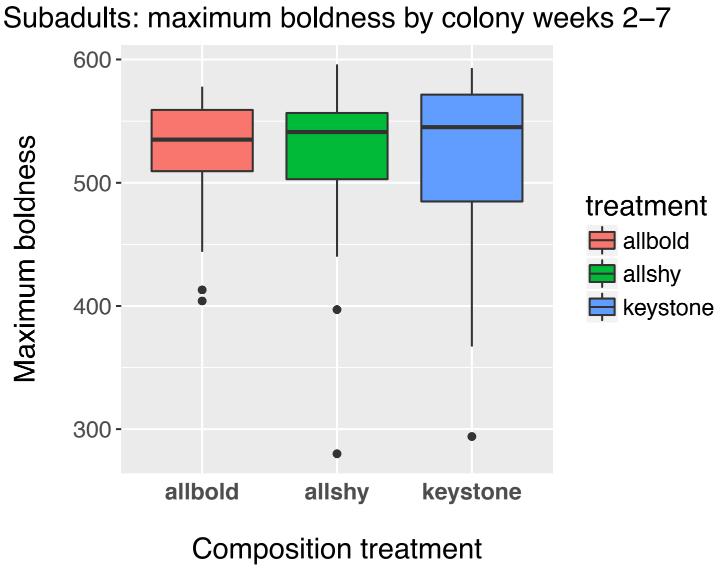


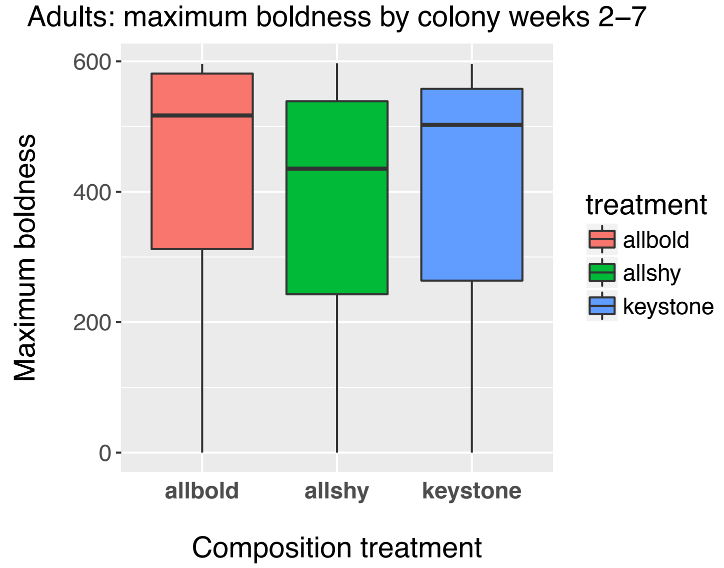


**Figure S4.** The distribution of maximum (keystone) boldness by initial composition treatment, excluding the first week when the artificial distributions were first created. The distributions are similar, indicating that keystone boldness became similar regardless of initial conditions.

**Table S1. Convergence of boldness distributions after first week.**

P values resulting from two-sample Kolmogorov-Smirnov test ('ks.test' in R 'stats' package), to compare the boldness distributions in the 3 different initial boldness treatments. $P < 0.05$ is evidence that the distributions are different.

In weeks 2-6 the distributions are substantially converged, except for 2 cases in both adults and subadults, though the p value remains relatively large for this test (> 0.001). It should also be noted that with multiple comparisons the expectation for a significant p value is raised.

**Adult spiders**

| Week | allbold vs allshy | allbold vs keystone | allshy vs keystone |
|---|---|---|---|
| 1 | 0.000 | 0.000 | 0.621 |
| 2 | 0.723 | 1.000 | 0.293 |
| 3 | 0.001 | 0.010 | 0.764 |
| 4 | 0.997 | 1.000 | 0.999 |
| 5 | 1.000 | 1.000 | 0.999 |
| 6 | 0.676 | 0.402 | 0.848 |

**Subadult spiders**

| Week | allbold vs allshy | allbold vs keystone | allshy vs keystone |
|---|---|---|---|
| 1 | 0.000 | 0.000 | 0.976 |
| 2 | 0.652 | 0.493 | 0.884 |
| 3 | 0.949 | 0.012 | 0.063 |
| 4 | 0.827 | 1.000 | 0.904 |
| 5 | 0.013 | 0.201 | 0.479 |
| 6 | 0.995 | 0.618 | 0.848 |

*Results of the separate adult and subadult spider model selection procedures*

**Adult spiders**

**Table S2.** Adults: Output of selected model for prey attack speed. In the fixed effect source colony, Col_num2, 3, measure the effect relative to baseline Col_num1. To measure the random effect of group and time, $\mu_i$ is the individual-specific time invariant random effect and $\nu_{it}$ is the individual-specific time varying residual random effect. 'Days before' is before prey stimulus, and is a fixed effect interacting with the chosen predictor variables.

| | Adult spiders: selected model for latency to attack | | | | | | |
|---|---|---|---|---|---|---|---|
| Level of Analysis | Coefficient | Days before | Estimate log(seconds) | Standard error | t value | p | Sig. at 5% |
| | (Intercept) | | 5.702 | 0.397 | 14.357 | < 0.001 | * |
| Individual | Closeness of keystone | 4 | 10.715 | 6.682 | 1.604 | 0.109 | |
| | | 2 | 12.766 | 6.269 | 2.036 | 0.042 | * |
| | | 0 | 18.842 | 7.205 | 2.615 | 0.0089 | * |
| Group | Average degree | 4 | -0.172 | 0.129 | -1.334 | 0.182 | |
| | | 2 | -0.239 | 0.150 | -1.596 | 0.111 | |
| | | 0 | -0.367 | 0.175 | -2.101 | 0.036 | * |
| Source colony (fixed effect) | Col_num2 | | 0.951 | 0.509 | 1.870 | 0.061 | |
| | Col_num3 | | -0.538 | 0.592 | -0.909 | 0.363 | |
| Group and time (random) | log(Sigma($\mu_i$)) | | 0.0187 | 0.192 | 0.097 | 0.923 | |
| | log(Sigma($\nu_{it}$)) | | 0.341 | 0.052 | 6.540 | < 0.001 | * |

**Table S3.** Generalized Variance Inflation Factors (VIFs) for the adult model. Calculated using the vif() function in the 'car' R package; the corrected GVIF takes into account categorical variables. The predictor variables maximum boldness and degree of keystone interact with measurement day (4, 2, or 0 days before prey attack assay).

| Coefficient | Df | GVIF^(1/(2*Df) |
|---|---|---|
| Source colony (Col_num) | 2 | 1.006 |
| Closeness of keystone : day | 3 | 1.717 |
| Average degree : day | 3 | 1.716 |

The corrected GVIFs for maximum boldness and degree of keystone are not of concern (squared corrected GVIF < 10, also see (O'Brien, 2007)).

**Subadult spiders**

**Table S4.** Subadults: Output of selected model for prey attack speed. Boldness is measured on a 0-600 scale. In the fixed effect source colony, Col_num2, 3, 4 measure the effect relative to baseline Col_num1. To measure the random effect of group and time, $\mu_i$ is the individual-specific time invariant random effect and $\nu_{it}$ is the individual-specific time varying residual random effect.

| | Adult spiders: selected model for latency to attack | | | | | | |
|---|---|---|---|---|---|---|---|
| Level of Analysis | Coefficient | Days before | Estimate log(seconds) | Standard error | t value | p | Sig. at 5% |
| | (Intercept) | | 7.262 | 0.965 | 7.523 | < 0.001 | * |
| Individual | Maximum boldness | 4 | -0.0053 | 0.0015 | -3.548 | < 0.001 | * |
| | | 2 | -0.0049 | 0.0016 | -3.045 | 0.002 | * |
| | | 0 | -0.0051 | 0.0016 | -3.210 | 0.001 | * |
| Group | Average degree | 4 | 0.0052 | 0.0247 | 0.211 | 0.833 | |
| | | 2 | -0.0410 | 0.0031 | -1.304 | 0.192 | |
| | | 0 | -0.0245 | 0.0245 | -1.000 | 0.317 | |
| Source colony (fixed effect) | Col_num2 | | 1.935 | 0.748 | 2.587 | 0.009 | * |
| | Col_num3 | | -0.078 | 0.798 | -0.097 | 0.923 | |
| | Col_num4 | | -0.212 | 0.696 | -0.305 | 0.761 | |
| Group and time (random) | log(Sigma($\mu_i$)) | | -0.104 | 0.225 | -0.462 | 0.644 | |
| | log(Sigma($\nu_{it}$)) | | 0.149 | 0.059 | 2.529 | 0.011 | * |

**Table S5.** Generalized Variance Inflation Factors (VIFs) for subadult model. Calculated using the vif() function in the 'car' R package; the corrected GVIF takes into account categorical variables. The predictor variables maximum boldness and degree of keystone interact with measurement day (4, 2, or 0 days before prey attack assay).

| Coefficient | Df | GVIF^(1/(2*Df) |
|---|---|---|
| Source colony (Col_num) | 3 | 1.012 |
| Maximum boldness : day | 3 | 1.565 |
| Average degree : day | 3 | 1.564 |

The corrected GVIFs for maximum boldness and degree of keystone are not of concern (squared corrected GVIF < 10, also see (O'Brien, 2007)).

**Table S6.** An alternative subadult model with degree of keystone as a second individual-level effect, instead of the group-level effect average degree (Table S4). Maximum boldness remains the only significant predictor.

| | Adult spiders: selected model for latency to attack | | | | | | |
|---|---|---|---|---|---|---|---|
| Level of Analysis | Coefficient | Days before | Estimate log(seconds) | Standard error | t value | p | Sig. at 5% |
| | (Intercept) | | 6.858 | 1.0045 | 6.827 | < 0.001 | * |
| Individual | Degree of keystone | 4 | 0.032 | 0.019 | 1.709 | 0.087 | |
| | | 2 | 0.0029 | 0.022 | 0.131 | 0.896 | |
| | | 0 | 0.023 | 0.020 | 1.180 | 0.238 | |
| | Maximum boldness | 4 | -0.0050 | 0.0015 | -3.378 | < 0.001 | * |
| | | 2 | -0.0049 | 0.0016 | -3.040 | 0.002 | * |
| | | 0 | -0.0053 | 0.0016 | -3.374 | < 0.001 | * |
| Source colony (fixed effect) | Col_num2 | | 2.093 | 0.834 | 2.511 | 0.012 | * |
| | Col_num3 | | -0.018 | 0.807 | -0.022 | 0.983 | |
| | Col_num4 | | -0.322 | 0.763 | -0.422 | 0.673 | |
| Group and time (random) | log(Sigma($\mu_i$)) | | -0.040 | 0.212 | -0.189 | 0.850 | |
| | log(Sigma($\nu_{it}$)) | | 0.143 | 0.0587 | 2.440 | 0.015 | * |

| Coefficient | Df | GVIF^(1/(2*Df) |
|---|---|---|
| Source colony (Col_num) | 3 | 1.012 |
| Degree of keystone : day | 3 | 1.384 |
| Maximum boldness : day | 3 | 1.393 |

**Table S7. Adult spiders – Full-model Averaging Results**

Parameter and variance estimates are made according to

$$\tilde{\bar{\beta}} = \sum_{i=1}^{R} w_i \hat{\beta}_i \qquad \widehat{var}\left(\tilde{\bar{\beta}}\right) = \sum w_i \left[\widehat{var}\left(\hat{\beta}_i\right) + \left(\beta_i - \tilde{\bar{\beta}}\right)^2\right]$$

Where $w_i$ is the Akaike model weight. See (Symonds and Moussalli, 2011) and (Lukacs et al. 2009).

For the adult spiders, none of the predictors are significant under full-model averaging, suggesting that excessive model selection uncertainty makes this approach uninformative. Instead, a parsimonious model selected using predictor ranking (Table S2) allows us to identify informative predictors.

| | Adult spiders: Full-model Averaging Results | | | | | |
|---|---|---|---|---|---|---|
| Level of Analysis | Coefficient | Days before | Estimate log(seconds) | Standard error | t value | Sig. at 5% |
| | (Intercept) | | 5.6852 | 0.6774 | 8.3925 | * |
| Individual | Degree of keystone | 4 | -0.0147 | 0.0540 | -0.2717 | |
| | | 2 | -0.0095 | 0.0520 | -0.1830 | |
| | | 0 | -0.0147 | 0.0623 | -0.2354 | |
| | Keystone closeness | 4 | 8.4980 | 6.9952 | 1.2148 | |
| | | 2 | 8.3614 | 6.7450 | 1.2397 | |
| | | 0 | 12.0574 | 8.8400 | 1.3640 | |
| | Maximum boldness | 4 | 0.0003 | 0.0006 | 0.5155 | |
| | | 2 | 0.0004 | 0.0008 | 0.5032 | |
| | | 0 | 0.0003 | 0.0007 | 0.4118 | |
| Subgroup | Modularity | 4 | -0.5745 | 0.9094 | -0.6317 | |
| | | 2 | -0.5248 | 0.8842 | -0.5935 | |
| | | 0 | -0.4508 | 0.7923 | -0.5690 | |
| Group | Average degree | 4 | -0.0896 | 0.1386 | -0.6463 | |
| | | 2 | -0.1302 | 0.1859 | -0.7006 | |
| | | 0 | -0.1828 | 0.2450 | -0.7464 | |
| | Skewness of degree distribution | 4 | 0.0011 | 0.0620 | 0.0171 | |
| | | 2 | -0.0011 | 0.0580 | -0.0190 | |
| | | 0 | -0.0057 | 0.0773 | -0.0744 | |
| Source colony (fixed effect) | Col_num2 | | 0.9687 | 0.5320 | 1.8208 | |
| | Col_num3 | | -0.5438 | 0.6104 | -0.8910 | |
| Group and time (random) | log(Sigma($\mu_i$)) | | 0.0437 | 0.1926 | 0.2267 | |
| | log(Sigma($\nu_{it}$)) | | 0.3416 | 0.0533 | 6.4119 | * |

**Table S8. Subadult spiders – Full-model Averaging Results**

Calculated with the equations given at Table S7. The same qualitative result is found as in Table S4: maximum boldness is a significant predictor of attack speed.

| | Subadult spiders: Full-model Averaging Results | | | | | |
|---|---|---|---|---|---|---|
| Level of Analysis | Coefficient | Days before | Estimate log(seconds) | Standard error | t value | Sig. at 5% |
| | (Intercept) | | 7.0716 | 1.0905 | 6.4847 | * |
| Individual | Degree of keystone | 4 | 0.0281 | 0.0344 | 0.8158 | |
| | | 2 | 0.0118 | 0.0231 | 0.5117 | |
| | | 0 | 0.0247 | 0.0324 | 0.7643 | |
| | Keystone closeness | 4 | 0.8071 | 2.8725 | 0.2810 | |
| | | 2 | 0.9217 | 3.1178 | 0.2956 | |
| | | 0 | 0.2378 | 2.2045 | 0.1078 | |
| | Maximum boldness | 4 | -0.0050 | 0.0017 | -2.9086 | * |
| | | 2 | -0.0053 | 0.0019 | -2.7946 | * |
| | | 0 | -0.0049 | 0.0018 | -2.7588 | * |
| Subgroup | Modularity | 4 | -0.1258 | 0.5981 | -0.2103 | |
| | | 2 | 0.7506 | 1.1170 | 0.6720 | |
| | | 0 | -0.5798 | 0.9804 | -0.5914 | |
| Group | Average degree | 4 | -0.0216 | 0.0404 | -0.5347 | |
| | | 2 | -0.0304 | 0.0493 | -0.6162 | |
| | | 0 | -0.0366 | 0.0477 | -0.7661 | |
| | Skewness of degree distribution | 4 | 0.0056 | 0.0819 | 0.0681 | |
| | | 2 | -0.0493 | 0.1465 | -0.3362 | |
| | | 0 | -0.0133 | 0.0727 | -0.1835 | |
| Source colony (fixed effect) | Col_num2 | | 2.0068 | 0.7807 | 2.5704 | * |
| | Col_num3 | | -0.0197 | 0.7960 | -0.0247 | |
| | Col_num4 | | -0.2374 | 0.7411 | -0.3203 | |
| Group and time (random) | log(Sigma($\mu_i$)) | | -0.0853 | 0.2191 | -0.3894 | |
| | log(Sigma($\nu_{it}$)) | | 0.1335 | 0.0612 | 2.1823 | * |
